# Genetic coupling of life-history and aerobic performance in Atlantic salmon

**DOI:** 10.1101/2021.08.23.457324

**Authors:** Jenni M. Prokkola, Eirik R. Åsheim, Sergey Morozov, Paul Bangura, Jaakko Erkinaro, Annukka Ruokolainen, Craig R. Primmer, Tutku Aykanat

## Abstract

A better understanding of the genetic and phenotypic architecture underlying life-history variation is a longstanding aim in biology. Theories suggest energy metabolism determines life-history variation by modulating resource acquisition and allocation trade-offs, but the genetic underpinnings of the relationship and its dependence on ecological conditions have rarely been demonstrated. The strong genetic determination of age-at-maturity by two unlinked genomic regions (*vgll3* and *six6*) makes Atlantic salmon (*Salmo salar*) an ideal model to address these questions. Using more than 250 juveniles in common garden conditions, we quantified the covariation between metabolic phenotypes –standard and maximum metabolic rates (SMR and MMR), and aerobic scope (AS) – and the life-history genomic regions and tested if food availability modulates the relationships. We found that the early maturation genotype in *vgll3* was associated with higher MMR and consequently AS. Additionally, MMR exhibited physiological epistasis; it was decreased when late maturation genotypes co-occurred in both genomic regions. Contrary to our expectation, the life-history genotypes had no effects on SMR. Further, food availability had no effect on the genetic covariation, suggesting a lack of genotype-by-environment interactions. Our results provide insights on the key organismal processes that link energy use at the juvenile-stage to age-at-maturity, indicating potential mechanisms by which metabolism and life-history can coevolve.

## Introduction

Physiological processes control how life-history diversity emerges from resource allocation and acquisition trade-offs^1^. The rate of aerobic energy metabolism is a pivotal mechanism contributing to life-history variation – it modulates resource acquisition, provides cells with ATP, and constrains energy allocation to different body components and functions. Theories, such as the metabolic theory of ecology and the pace-of-life syndrome theory^2,3^, suggest metabolic rate covaries with life-history variation within and among species. This covariation may have a genetic basis, consequently constraining trait evolution^4^, yet only a few studies have demonstrated intraspecific genetic covariation or co-evolution between metabolic rate and life-history traits^5-7^. Determining whether this relation is modified by different ecological contexts (e.g., food availability) is crucial to better understand the mechanisms shaping life history variation and demographic shifts in populations in response to environmental changes^8^.

The quintessential components of energy metabolism at the organismal level (i.e., the metabolic phenotypes) are standard metabolic rate (SMR), maximum metabolic rate (MMR), and the absolute aerobic scope (AS) that is the difference between SMR and MMR^9-11^. SMR is the minimum metabolic rate of an ectothermic animal associated with self-maintenance, and therefore defines the minimal cost of living (excluding growth, digestion, and locomotion). MMR defines the upper limit of aerobic performance that is functionally linked to SMR and the capacity to increase oxygen uptake and delivery beyond SMR. SMR and MMR together are the integral components of AS, which is the measure of surplus energy that can be allocated into non-maintenance functions, such as locomotion and digestion^12^. Higher AS is predicted to increase fitness via facilitating energetically demanding behaviours (such as migration, aggression, predator avoidance, and prey capture) and tolerance to environmental stress^11,13-15^. However, high aerobic performance comes with costs, including maintaining a larger heart and gill surface area (associated with increased demand for osmoregulation)^12,16,17^.

Allocation of energy to growth or improved condition can link metabolic phenotypes to life-history traits^18^. Life-history traits, such as the timing of maturation and migration, are determined by adaptive body-size thresholds^19-21^, and metabolic phenotypes are often correlated with growth rate, albeit in a context-dependent manner^11^. Under high food availability, a high SMR in combination with high AS can increase growth rate^22^, as it often correlates with traits that improve resource acquisition, such as dominance and digestive capacity^23-25^. Under low food availability, the growth benefit of high SMR or AS can be minimized (or even reversed for SMR) due to high self-maintenance costs^22,26,27^. In addition, individuals can be forced to seek new habitats or take more risks to acquire resources, exerting further fitness costs^28,29^. Hence, covariation between life-history and metabolism, whether it is genetically or environmentally driven, could be modulated by resource availability. A resource dependent change in genetic covariation, i.e., genotype-by-environment interaction, could maintain genetic variation in these traits^30-32^, but this has not been demonstrated.

In anadromous (sea-migrating) salmonids, the number of years the fish spends at sea before the first spawning, i.e., sea age-at-maturity, has a dramatic effect on its size-at-maturity^33^. Individuals spending one year at sea typically weigh 1–3 kg compared with 10–20 kg after three or more years. Increased size in late maturing individuals also translates to marked gains in reproductive investment in both sexes^33^. Earlier maturation, i.e., less time spent at sea, provides a potential fitness advantage through a higher probability of survival prior to reproduction and a shorter generation time, but comes at the expense of decreased fecundity and mating success (due to smaller size at reproduction^34-36^. The probability of early maturation at sea is also positively associated with faster growth and fat deposition in the freshwater (juvenile) stage, since the size of salmon at the onset of sea-migration has a significant influence on maturation timing at sea^19,37-40^. These relationships suggest that early maturation in salmon may be associated with higher SMR or aerobic scope via resource utilization already in early life-stages prior to sea migration^24,25^. In addition, metabolic phenotypes in the juvenile stage may explain maturation at sea through genetic correlations across life stages (e.g.,^41^) if metabolic rate or aerobic performance at sea is linked to earlier maturation.

In Atlantic salmon (*Salmo salar* L. 1758), a large proportion of variation in age-at-maturity is explained by a single genomic region that encompasses the *vgll3* gene on chromosome 25 42,43. In addition, variation in another locus on chromosome 9, *six6*, is a strong predictor of mean age-at-maturity among populations ^43^ and associated with early maturation in aquaculture salmon^44^. *Vgll3* and *six6* are also associated with size-at-maturity, with the alleles conferring late maturation being associated with larger age-specific body size especially after multiple years at sea^43^. Moreover, *vgll3* is associated with precocious maturation in male salmon parr via body condition^45^, emphasizing the causal effect of energy acquisition on maturation. In the last few decades, many Atlantic salmon populations have been maturing, on average, at younger ages^46^, which is associated with a change in *vgll3* allele frequency in some cases^47^. Recently, a link was found between the decrease in salmon age-at-maturity and a change in prey species composition^48^ (see also ^49^ for diet composition in relation to *six6*). These observations further highlight that genotype dependent differences in SMR or aerobic performance, which can contribute to foraging success or food assimilation (e.g.,^22,50,51^), may be related to contemporary life-history evolution in Atlantic salmon.

The strong effects of the *six6* and *vgll3* genomic regions on life-history variation provide an opportunity for the genetic covariation between age-at-maturity and energy metabolism to be studied prior to maturation, i.e., at the juvenile stage, by genetic prediction. This approach makes controlled, empirical settings more feasible, as salmon require several years to reach maturation. In this study, we test if genetic covariation exists between life-history and metabolic phenotypes in juveniles; we expect that early maturation genotypes show a higher metabolic activity (SMR, MMR, and AS) than late maturation genotypes under high food availability (e.g.,^7^). Further, because low resource availability weakens the relationship between SMR and growth^27^ and may constrain aerobic performance^8^, we also explore if food availability modulates the genetic covariation.

## Material and Methods

The experiments were conducted under an animal experiment permit granted by the Finnish Project Authorisation Board (permit nr. ESAVI/4511/2020). Additional experimental details are provided in online supplemental material.

### Fish rearing and genotyping

The parental Atlantic salmon (Table S1) were first-generation hatchery brood stock (from the river Kymijoki in Finland) from Laukaa hatchery, managed by the Natural Resources Institute Finland (LUKE). In October 2019, eggs and milt were transferred to the University of Helsinki for fertilisation. Full-sib families were crossed using heterozygous parents in *vgll3* and *six6* loci, (*vgll3*\*E/L and *six6*\*E/L where E and L refer to the alleles associated with early and late maturation, respectively). This provides offspring with all genotype combinations within each full-sib family. Feed rations were calculated assuming feed conversion efficiency of 0.8, using growth predictions by Elliott and Hurley^52^. In July 2020, fish were tagged with passive integrated transponder (PIT) tags and genotyped from fin clips.

### Experimental design

Food treatments were started in August 2020. Briefly, fish were fed twice week^-1^ in the low food treatment (Fig. 1). This was preferred over constant low ration to minimize dominance hierarchies in tanks^53^. Fish in the high food treatment were fed with the total ad-libitum daily ration. After 28–31d in the treatments, 48 and 32 fish from each family in the low and high food treatment, respectively (192 and 128 in total), were measured for their SMR and MMR (Fig. 1), though only 290 homozygous individuals were used in the analysis (Table S2).

**Fig. 1.**
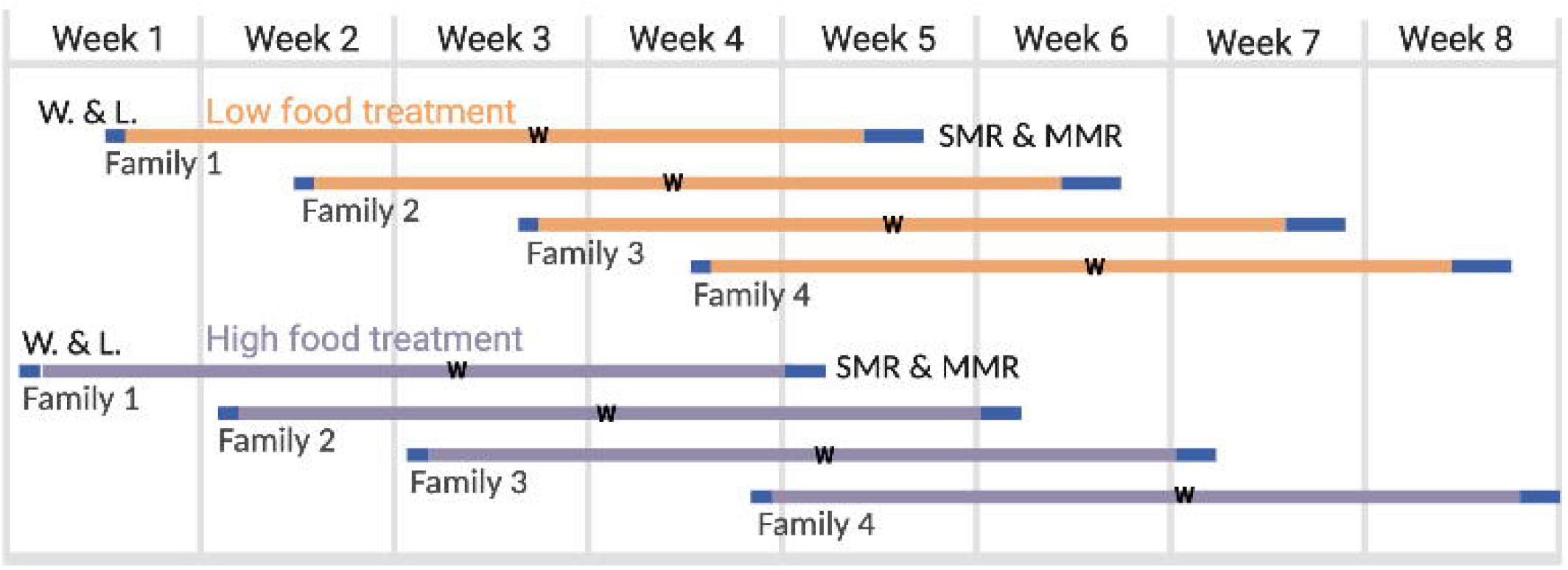
Timeline of the experiment. The duration of low food and high food treatments was four weeks. “W” indicates when fish density of high food tanks was reduced to the level of low food tanks. Each horizontal line represents a separate tank. Blue blocks represent procedures (W. & L. = weight and length, and/or SMR & MMR measurements).

### SMR and MMR measurements

Sixteen fish at once were moved into an acclimation tank two days before SMR measurement (Fig. S3). Each batch was from the same family and tank and balanced for genotype-sex-combinations. During acclimation, the fish were held individually, without feeding at 11°C ± 0.1°C in 20×20×10cm cages, to minimize the effects of temperature fluctuations, digestion, growth, and social interactions on SMR^54^. We measured SMR using intermittent flow respirometry^55-57^. The SMR measurements commenced after 42–47h acclimation, between 11:30 and 14:30, until 8:00 the following day. Afterwards, MMR was measured using the chase method, similarly to, e.g.,^58^, where MMR reflects increased aerobic respiration related to exercise and the oxygen debt incurred by anaerobic respiration ^59,60^. Fish were then euthanized with an overdose of methanesulfonate, measured, and weighed. All fish were confirmed to have immature gonads.

### Analysis of respirometry data

For SMR, oxygen consumption rate (*MO*_2_, mg O_2_ h^-1^) for each linear measurement phase was derived from best-fit linear regression of dissolved oxygen concentration over time. The mean of the lowest normal distribution (MLND) was used to estimate SMR for each individual as mg O_2_ h^-1^ from the extracted *MO*_2_ slopes^10^.

MMR was calculated for each individual as mg O_2_ h^-1^ from the O_2_ concentrations after performing background correction in *FishResp*^61^. We used two methods to identify the slope of the steepest decrease in O_2_ saturation. First, we used *respR*^62^ with the function *auto_rate*, fitting one and two-minute windows^63^ (example slope in Fig. S5A). Second, slopes for MMR were extracted using a derivative of a polynomial curve fitted on each measurement (function *smooth*.*spline*, df=10). This is the ‘*spline-MMR*’ method (Fig. S5B). The slopes were then used to calculate MMR in mg O_2_ h^-1^ using *FishResp*-package function *calculate*.*MR*. MMR values calculated by the 1-min *respR* and *spline-MMR* approaches were highly correlated (Pearson-r = 0.98, 95% confidence interval 0.98 – 0.99). We selected the spline-MMR data for further analysis.

### Statistical analyses

Data were analysed in R v. 3.6.2^64^. To test for the effects of treatment and genotype on metabolic variables, we ran separate linear mixed models using SMR, MMR, and AS as response variables. The response variables and body mass were log_10_-transformed to account for allometric scaling of metabolic rate, fixed effects were centred, and covariates were centred and scaled^65^ (Table S3). We included treatment, *vgll3* and *six6* genotypes, and sex as fixed effects in all models. For SMR, family and measurement batch were used as random terms (models including chamber as random term were singular, and no variance was explained by chamber). For MMR and AS, family, person performing the chase test, and chamber identity were included as random terms. The order in which pairs of fish were tested for MMR each day (values 1–8) was included as a covariate for MMR and AS. To test if genotype-specific metabolic rates were affected by sex and food availability, we fitted pairwise interactions between *vgll3* and *six6* genotypes, between genotypes and treatment, and between genotypes and sex. The interaction of log10 body mass with treatment was included to test for potential treatment-specific allometric scaling of metabolic rate.

The full models were fitted using *lme4* v. 1.1-26 ^66^ with an alpha value 0.05. P- and F-test values for fixed effects were computed using type III tests with Satterthwaite’s method. Pairwise differences between significant interaction terms were obtained by post hoc analysis using *emmeans*^67^. Residuals of models were confirmed to be homoscedastic and normally distributed (but the residuals of SMR between the high food and low food treatments were slightly heteroscedastic). One outlier was identified and removed from each of SMR and MMR (using *outlierTest* in package *car*, Bonferroni-corrected p < 0.05). The proportion of variance explained by genotypes was calculated with *partR2*^68^. Predicted means were obtained with *ggpredict* in package *ggeffects*^*69*^. The data were visualized using *ggplot2* v.3.3.370 and *interactions*^*71*^. Pearson’s correlation coefficient among mass- and family-corrected SMR, MMR and AS were calculated using residuals from a mixed model.

We also evaluated alternative models, and assess parameter significance using the corrected Akaike information criterion score (AICc), an AIC score with a stronger penalty for complex models^72^, using the *dredge* function from package *MuMIn*^73^. We employed model averaging if more than one model was similarly parsimonious (Table S4), in which parameter estimates were obtained from weighted averages of all best models^74^ (i.e., models with ΔAICc <2 compared to the most parsimonious models) using *model*.*avg* function (subset option = full) from package *MuMIn*^73^.

## Results

Low food treatment decreased both the specific growth rate and condition factor of the fish compared to high food treatment (Fig. S6). The mean body length of fish was 70.6 ± 4.5 and 66.2 ± 4.9 mm (SD), and the mean body mass was 4.2 ± 0.8 and 3.3 ± 0.8g after the high and low food treatment, respectively.

### Standard metabolic rate

There was no significant genotype, food availability or sex effect on SMR (Table 1, Fig. 2A). There was a marginally significant interaction effect of *six6* and food availability on SMR in the full model (p = 0.045) but this was non-significant in the averaged model (Table S5), and none of the pairwise contrasts were significant (the largest effect being: *six6* EE-genotype, high food vs. low food, t_25.6_ = −2.37, p = 0.11). The metabolic scaling exponent, *b*, i.e., the slope of log SMR with log body mass, was marginally higher in the high food treatment (0.94, R^2^ = 0.73) than in the low food treatment (0.87, R^2^ = 0.69, Fig. S7a, p = 0.047 (Table 1), p = 0.08 (Table S5)).

**Table 1.** Linear mixed models for log_10_-transformed metabolic phenotypes. All variables were centred to a mean of 0 (the category with a positive value is shown in parentheses), log_10_ body mass was scaled and centred. Significant effects shown in bold.

|  | Fixed effect | Estimate | SE | SSq | Den<br>DF | F | p | Random<br>effect | Var (95% CI) |
| --- | --- | --- | --- | --- | --- | --- | --- | --- | --- |
| SMR | Intercept | -0.295 | 0.019 |  |  |  |  |  |  |
|  | Treatment (LF) | 0.019 | 0.012 | 0.0039 | 17.67 | 2.772 | 0.114 |  |  |
|  | Sex (male) | 0.002 | 0.005 | 0.0002 | 248.11 | 0.172 | 0.679 |  |  |
|  | *Vgll3* (LL) | -0.002 | 0.005 | 0.0003 | 248.98 | 0.235 | 0.628 |  |  |
|  | *Six6* (LL) | 0.002 | 0.005 | 0.0002 | 248.11 | 0.172 | 0.679 |  |  |
|  | **Log10 BM** | **0.102** | **0.003** | **1.4570** | **259.04** | **1025.739** | **<0.0001** |  |  |
|  | **Treatment (LF):log10 BM** | **0.013** | **0.006** | **0.0057** | **259.25** | **3.987** | **0.047** |  |  |
|  | Treatment (LF):*Vgll3* (LL) | 0.006 | 0.010 | 0.0006 | 249.25 | 0.410 | 0.523 |  |  |
|  | **Treatment (LF):*Six6* (LL)** | **-0.020** | **0.010** | **0.0058** | **249.21** | **4.080** | **0.044** |  |  |
|  | Sex (male):*Vgll3* (LL) | 0.001 | 0.010 | 0.00001 | 248.33 | 0.004 | 0.948 | Batch | 0.0005 (0.0002, 0.0012) |
|  | Sex (male):*Six6* (LL) | 0.001 | 0.009 | 0.00001 | 247.62 | 0.006 | 0.937 | Family | 0.0013 (0.0003, 0.0083) |
|  | *Vgll3* (LL):*Six6* (LL) | -0.016 | 0.009 | 0.0042 | 248.20 | 2.930 | 0.088 | Residual | 0.0014 |
| MMR | Intercept | 0.333 | 0.013 |  |  |  |  |  |  |
|  | Treatment (LF) | 0.008 | 0.006 | 0.003 | 249.48 | 1.632 | 0.203 |  |  |
|  | Sex (male) | -0.003 | 0.005 | 0.001 | 254.25 | 0.295 | 0.588 |  |  |
|  | ***Vgll3* (LL)** | **-0.017** | **0.005** | **0.017** | **252.06** | **10.429** | **0.001** |  |  |
|  | *Six6* (LL) | -0.006 | 0.005 | 0.002 | 252.45 | 1.371 | 0.243 |  |  |
|  | **Log10 BM** | **0.088** | **0.003** | **1.231** | **256.37** | **737.850** | **<0.0001** |  |  |
|  | Test order | -0.003 | 0.004 | 0.001 | 13.81 | 0.769 | 0.396 |  |  |
|  | Treatment (LF):log10 BM | -0.010 | 0.007 | 0.004 | 255.47 | 2.252 | 0.135 |  |  |
|  | Treatment (LF):*Vgll3* (LL) | -0.011 | 0.011 | 0.002 | 253.11 | 1.057 | 0.305 |  |  |
|  | Treatment (LF):*Six6* (LL) | -0.010 | 0.011 | 0.001 | 255.77 | 0.840 | 0.360 | Chamber | 0.0002 (0, 0.0005) |
|  | Sex (male):*Vgll3* (LL) | 0.016 | 0.010 | 0.004 | 249.17 | 2.474 | 0.117 | Initial | 0.0001 (0, 0.0012) |
|  | Sex (male):*Six6* (LL) | 0.008 | 0.010 | 0.001 | 251.83 | 0.654 | 0.420 | Family | 0.0004 (0.0001, 0.0027) |
|  | ***Vgll3* (LL) x *Six6* (LL)** | **-0.023** | **0.010** | **0.008** | **253.67** | **5.076** | **0.025** | Residual | 0.0017 |
| AS | Intercept | 0.210 | 0.019 |  |  |  |  |  |  |
|  | Treatment (LF) | 0.012 | 0.008 | 0.006 | 238.60 | 2.168 | 0.142 |  |  |
|  | Sex (male) | -0.004 | 0.007 | 0.001 | 243.05 | 0.443 | 0.507 |  |  |
|  | ***Vgll3* (LL)** | **-0.016** | **0.007** | **0.015** | **240.86** | **5.653** | **0.018** |  |  |
|  | *Six6* (LL) | -0.008 | 0.007 | 0.004 | 242.39 | 1.468 | 0.227 |  |  |
|  | **Log10 BM** | **0.089** | **0.004** | **1.183** | **245.15** | **437.482** | **<0.0001** |  |  |
|  | Test order | -0.003 | 0.005 | 0.001 | 14.796 | 0.387 | 0.543 |  |  |
|  | **Treatment (LF):log10 BM** | **-0.018** | **0.009** | **0.012** | **245.12** | **4.409** | **0.037** |  |  |
|  | Treatment (LF):*Vgll3* (LL) | -0.016 | 0.014 | 0.003 | 241.59 | 1.231 | 0.268 |  |  |
|  | Treatment (LF):*Six6* (LL) | 0.001 | 0.014 | 0.000 | 244.70 | 0.002 | 0.964 | Chamber | 0.0002 (0, 0.0008) |
|  | Sex (male):*Vgll3* (LL) | 0.023 | 0.014 | 0.008 | 239.76 | 2.802 | 0.095 | Initial | 0.0002 (0, 0.0022) |
|  | Sex (male):*Six6* (LL) | 0.009 | 0.014 | 0.001 | 239.97 | 0.487 | 0.486 | Family | 0.001 (0.0003, 0.0064) |
|  | *Vgll3* (LL) x *Six6* (LL) | -0.020 | 0.014 | 0.006 | 243.02 | 2.086 | 0.150 | Residual | 0.0027 |

BM = Body mass, LF = Low food

**Fig 2.**
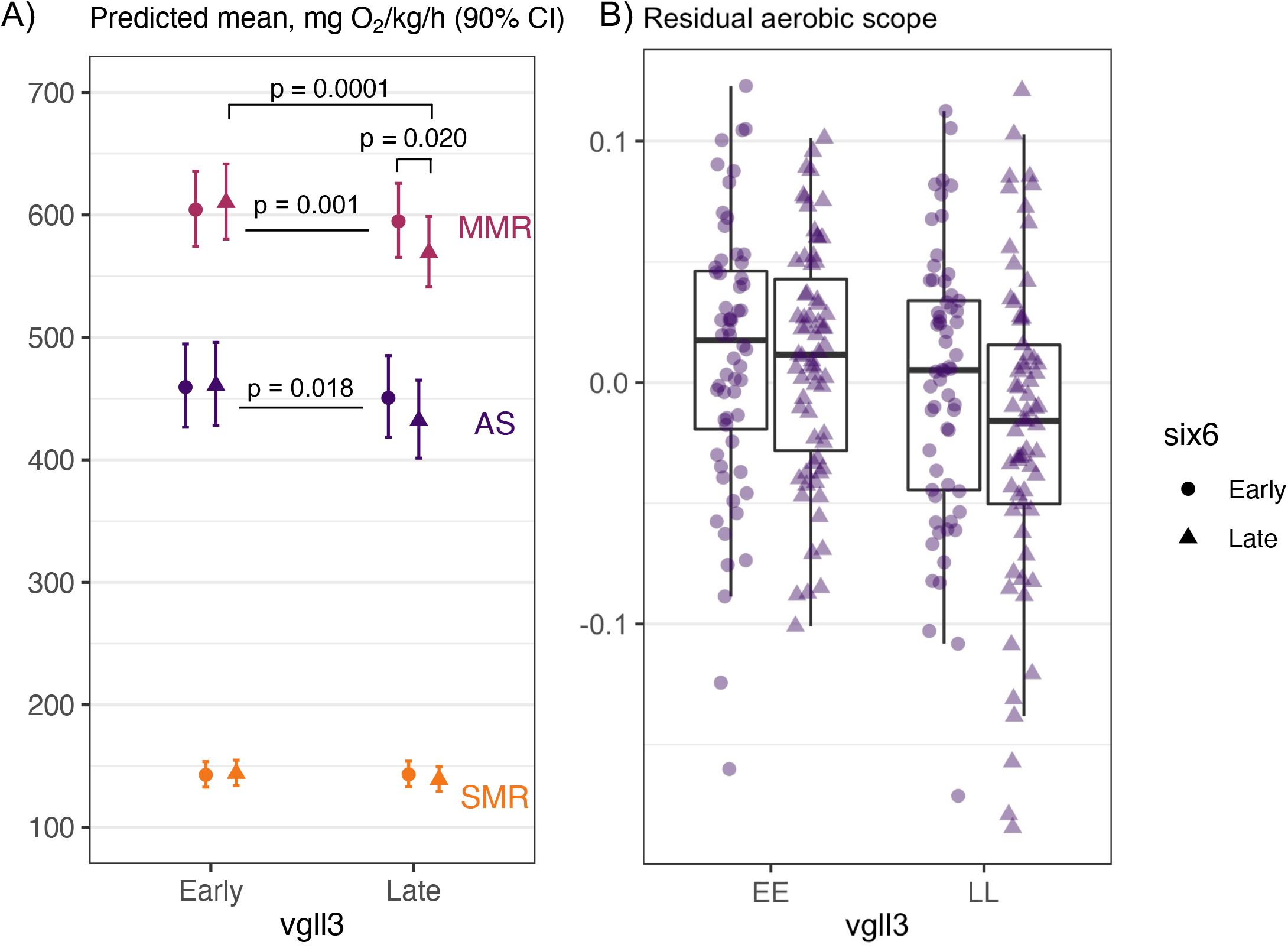
A) Predicted means for standard metabolic rate (SMR), maximum metabolic rate (MMR), and aerobic scope (AS) in *vgll3* and *six6* early- and late-maturation genotypes with 90% confidence intervals. The means are for average body mass, treatment, and sex effects, and back transformed to linear scale. P-values show significant pairwise differences between genotypes for MMR on top of the points, and for the *vgll3* main effects (Table 1) in MMR and AS between the points. N = 60–71 in each genotype combination (same individuals used for all traits). B) Residual aerobic scope from a linear mixed model including log10 aerobic scope as response, log10 body mass as predictor and family as random term, showing individuals (by points) in each genotype combination.

### Maximum metabolic rate

Fish with the *vgll3* early maturation genotype had a higher MMR than fish with the late maturation genotype (Fig. 2A, Table 1). *Vgll3* genotype also interacted with *six6*, such that MMR was decreased when late maturation genotypes of the two loci cooccurred compared to other genotype combinations (Fig. 2, Table 1). The genotype effects together explained ∼5% of the variance in MMR (Table S6). None of the treatment-genotype or sex-genotype interactions or the main effects of sex or food availability had a significant effect on MMR (Table 1, Table S5). Unlike SMR, the metabolic scaling of MMR was not significantly affected by food treatment (*b* = 0.86, R^2^ = 0.76).

### Aerobic scope

Fish with the *vgll3* early maturation genotype had a higher AS compared to the late maturation genotype (Fig. 2A-B, Table 1, predicted means 460.2 and 440.9 mg O_2_ kg^-1^ h^-1^ for early and late maturation genotypes, respectively). *Vgll3* was estimated to explain 1.7% of the variance in mass-corrected AS (Table S6). AS was marginally higher under low food availability than high food availability, but only in smaller fish (interaction p = 0.037 (Table 1) and p = 0.18 (Table S6)); scaling exponent *b* = 0.94 (R^2^ = 0.57) in the high food and 0.90 (R^2^ = 0.68) in the low food treatment (Fig. S7b). The *vgll3* and food treatment effects were also significant when mass adjusted SMR was included as a covariate in the model (Table S7), indicating that the genotype effect was independent of SMR. None of the other factors had a significant effect on AS (Table 1).

### Correlations among metabolic phenotypes

There was a positive correlation between mass- and family-corrected SMR and MMR in the high food, but not the low food treatment (Fig. S8), and a very strong correlation between MMR and AS in both treatments (Table 2).

**Table 2.** Pearson’s correlation coefficients between metabolic phenotypes in high food (above diagonal) and low food (below diagonal) treatments. P-values given in parentheses.

|  | rSMR | rMMR | rAbsAS |
| --- | --- | --- | --- |
| rSMR |  | **0.28 (0.004)** | 0.13 (0.09) |
| rMMR | 0.11 (0.16) |  | **0.99 (<0.001)** |
| rAbsAS | **-0.17 (0.04)** | **0.96 (<0.001)** |  |

## Discussion

The timing of maturation, just as many life-history traits, depends on reaching a certain body size threshold, i.e., the acquisition of sufficient energy that can be allocated for maturation processes^38,39,75,76^. In line with our hypothesis, we found that the *vgll3* early maturation genotype increased the aerobic scope (AS) of juvenile Atlantic salmon compared to the late maturation genotype. This effect was driven by a change in maximum metabolic rate (MMR), not standard metabolic rate (SMR) (nearly all variation in AS was explained by MMR in our study, Table 2). A previous study showed that higher condition factor, mediated by the *vgll3* early maturation genotype, positively affected the initiation of male maturation^45^. The results presented here suggest that superior resource acquisition or assimilation via higher AS, driven by a higher MMR, is a potential mechanism by which an increased condition factor in individuals with the *vgll3* early maturation genotype could be achieved compared conspecifics with the late maturation genotype. In addition to differences in mean performance between *vgll3* genotypes, the two loci in our study exhibited physiological epistasis^77^, as the cooccurrence of the late maturing genotypes in both loci was associated with lower MMR than their additive effects. The epistasis may help to maintain genetic variation under rapid adaptive responses^78,79^.

The functional pathways that may explain the epistasis and the main effect of *vgll3* are not well known. However, both *six6* and *vgll3* are expressed during development and have been implicated in the control of cell fate commitment and the hypothalamus-pituitary-gonad axis in salmon^80,81^. Hence, the epistatic interaction between two genomic regions may stem from, e.g., developmental canalisation^82^. Addressing the causal physiological and morphological mechanisms of the link between the genomic regions and aerobic performance can shed light into the mechanisms of life history evolution in salmon; the *vgll3* genomic region is the major genetic axis explaining variation in age-at-maturity in salmon^43^, and variation in the locus is spatially divergent among populations and under rapid adaptative evolution^43,47,83,84^.

Because of context-dependent covariation between metabolism and growth rate, whereby high SMR improves growth under high, but not low, resource availability^27^ and because aerobic performance can be reduced under food limitation (e.g.,^8^), we tested if food availability modified the genetic covariation between metabolic phenotypes and age-at-maturity. Against our predictions, there was no change in SMR or MMR due to feed restriction, nor did we find genotype-by-environment interactions, despite a strong decrease in growth rate in low food treatment. The results indicate that the different genotypes exhibit no plastic responses to food availability. However, a stronger feed deprivation, e.g., similar to those occurring in the winter^85^, could have induced a more pronounced effect on SMR^86,87^. Our low food treatment included approximately 3 days of fasting in between feeding to satiation, similar to a “feast and famine” feeding strategy^88^. A lack of metabolic response to reduced food availability may be beneficial if it allows the individual to maximize acquisition via food assimilation during “feasting”.

Unlike MMR (and consequently AS), SMR did not exhibit *vgll3*-linked covariation with age-at-maturity. A decoupling of SMR and MMR in relation to life-history variation was also found by Archer et al.^89^ in resident and migratory brown trout (*Salmo trutta*), and high MMR, but not SMR, was positively selected for in Atlantic salmon under high food competition in a study by Auer et al.^90^. Further, a lack of differences in SMR across the *vgll3* genotypes was found in our parallel study, in which the fish were smaller (mean ∼1g) compared to this study (mean ∼4g)^91^. The lack of association to the age-at-maturity loci is unexpected, as SMR, or basal metabolic rate in endotherms, has been proposed to explain life-history variation along the fast-slow axis^5,7^, but see^92^. Our results suggest that the genetic control of maturation by the *vgll3* genomic region via MMR mostly involves physiological pathways that do not alter SMR simultaneously. Such pathways may be related to oxygen demand by tissues or its supply during stress and/or exhaustive exercise. For example, structural and functional variation in the heart (i.e., cardiac output) or muscle, and mechanisms that modulate oxygen carrying capacity of the cardiovascular system might invoke changes in MMR without altering SMR^93-95^. However, our study does not rule out the possibility of the metabolic phenotypes affecting variation in age-at-maturity also phenotypically or via small-effect loci^44,96^.

The differences in aerobic performance between life-history genotypes may have arisen due to correlated selection mediated by resource acquisition: higher aerobic scope, which was associated with early maturation, could enable higher feeding capacity^50^ and improve foraging efficiency, for example, via a shorter searching time of prey^11,85^. Further, salmon in the wild are increasingly experiencing higher than optimal temperatures due to climate change^97^, and MMR is typically less plastic than SMR in response to environmental temperature^15^. Thus, our results indicate a potentially important advantage for individuals carrying the early maturation genotype under global warming, which could be mediated by a higher appetite^98^. This advantage may also extend to higher survival during spawning migration if the genotype effect on AS persists across life-stages^13,99,100^. It can also be relevant to survival of salmon after spawning, and thereby repeated spawning (iteroparity), which is co-inherited with the same *vgll3* genotype as early maturation^101^. However, the lack of sex differences in metabolic phenotypes in our study both across and within age-at-maturity genotypes, suggests that sex-dependent life-history variation in salmon^33^ is not reflected in metabolic rates during the juvenile stage (see also ^91^).

Although a causal relationship between metabolism at the juvenile stage and age at maturity would not be surprising, pleiotropy or linkage provide alternatives for the basis of the observed association. For example, the *vgll3* gene encodes a transcription cofactor associated with cell fate commitment and is expressed in many tissues^80,102^, thus it is likely pleiotropic with multiple independent functions. Similarly, several polymorphic loci with putative functional variation are co-localised (i.e., linked) in the genomic region^43^. Finally, stage-dependent genetic correlations in metabolism may obscure the time point that the trait influences the life-history variation. For instance, higher aerobic scope can induce maturation via facilitating the size attained both in the freshwater and at sea (e.g.,^19,39^), but whether aerobic scope genetically covaries across life stages is yet to be explored.

The presence of genetic covariation between aerobic scope at the juvenile stage and age-at-maturity at the *vgll3* genomic region suggests potential for multi-trait evolution across life-stages, whereby selection acting on either trait would alter the phenotypic variation of the other^4,103^. For example, if natural selection towards later age-at-maturity increases the frequency of late maturing allele, this would constrain the aerobic scope of juveniles in the population, even if that may be a suboptimal phenotype. On the other hand, genetic covariation may help to maintain optimal trait variation in age-at-maturity, i.e., by limiting potentially maladaptive environmentally induced (i.e., plastic) variation in age-at-maturity (e.g.,^104,105^). For example, river geophysical properties are important determinants of the optimal age structure at maturity, whereby populations in smaller tributaries have a younger, and populations in large, fast-flowing rivers have an older age structure^33^. Forecasting age-at-maturity from aerobic performance at earlier stages (e.g., via improved growth^22,51^) would result in maladaptive age structure if the covariation was explained entirely by environmental effects. However, our study was aimed at measuring the statistical association, and does not provide estimates using a classical quantitative genetic framework (i.e., we did not quantify environmental sources of variation, or variation due to technical or other genetic effects) or measure evolutionary change. Therefore, partitioning biologically meaningful covariation and quantifying the correlated response to selection were beyond the scope of this study^106^.

Understanding the physiological basis of life-history variation in different life-stages and environmental conditions can provide insights into the factors driving life-history evolution, and hence, better predictions of the responses of populations to environmental changes. Wild salmon populations have declined in recent decades, with a concomitant decrease in the frequency of late maturing individuals^47,97^. Our study used an eco-physiological approach to identify a potentially adaptive phenotype relating genetic variation and age-at-maturity in salmon and suggests that evolution towards an earlier age-at-maturity can cause correlated selection towards increased MMR and aerobic scope. In conclusion, the integration of age-at-maturity and aerobic performance in the early life-stages via simple genetic mechanisms, as shown in this study, could contribute to the diversification of ecotypes within species.

## Acknowledgements

We thank Nikolai Piavchenko, Katja Maamela, Markus Lauha, Petra Liljeström, Suvi Ikonen, Mikko Immonen, and Anna Toikkanen for fish care and technical help during the experiment, and Lammi Biological Station for a high-quality research environment. Dr. Markus Haapala and the Nordic University Hub NordicPOP (Nordforsk, project no. 85352) are acknowledged for the 3D printing service, and Heidrikur Bergsson for sharing the chamber cap model. The study was funded by Academy of Finland (T. Aykanat: 325964, 1328860, C. R. Primmer: 314254 and 314255, 327255), the European Research Council under the European Articles Union’s Horizon 2020 research and innovation program (grant no. 742312), and the University of Helsinki.

## Data availability

The data are available in https://doi.org/10.5281/zenodo.5667078. R codes for the analyses are available in an archived repository (v.1.0.2 https://doi.org/10.5281/zenodo.5783978).

## Supplemental Material

Pdf-file with Material and Methods, Figures, and Tables.

