## Supplemental material for "Genetic coupling of life-history and aerobic performance in Atlantic salmon"

Contents:

1. Material and Methods
2. Figures
3. Tables
4. References

### 1. Material and Methods

*Fish rearing and genotyping*

Eggs were incubated in vertical incubators at 7°C in the dark. On 6 March 2020, hatched alevins were transferred to Lammi Biological Station (Lammi, Finland) and released into circular tanks, diameter 90cm. Tank water volume was initially 189L and increased to 306L in July. Each family was reared in a separate tank, supplied with UV-filtered water from the nearby lake, Pääjärvi, warmed by 1°C with a heat-exchange system, under a simulated natural photoperiod. Feeding of alevins was started when most of the egg yolk had been consumed in March 2020.

Tanks were cleaned by scrubbing surfaces and siphoning excess food once week$^{-1}$ until June 1st, then approximately twice week$^{-1}$ until August 14th, then once week$^{-1}$ until the end of the experiment. Fish were initially fed with 0.2mm commercial feed (Vita, Veronesi), 4 times d$^{-1}$, then with 0.5mm feed and more frequent feeding, 16 times d$^{-1}$ using aquarium feeders. When the required feed weight exceeded 8g $d^{-1}$ we deployed Profi-Automatic feeders (LINN Gerätebau GmbH, Lennestad, Germany). Feeding was continued 'ad libitum', i.e., maximizing the number of feeds to up to 16 times $d^{-1}$. Mortality during March-July was approximately 8%. Water temperature during this time increased from *ca.* 4.5 °C to 11°C (Fig. S1). In July 2020, fish were anaesthetised with sodium bicarbonate-buffered methanesulfonate (100 mg/L) and individually tagged with passive integrated transponder (PIT) tags (length 8mm, width 1.2mm, Manruta, Quangdong, China) into the abdominal cavity, measured, and fin clipped (approx. 5% mortality during the procedure). After tagging, fish were released back to the tanks, where environmental enrichment was provided in the form of stones in three stainless-steel baskets.

DNA was extracted from fin clips and genotyped using Kompetitive Allele-Specific PCR (KASP, LGC genomics, UK; He *et al.* 2014) for *vgll3* (Sinclair-Waters *et al.* 2021), *six6*, and for genetic sex determination. The fin clips collected during PIT-tagging were placed in 20 µL of Lucigen QuickExtract DNA Extraction Solution 1.0 and kept on ice or at -20°C until extractions were completed on the day of sample collection according to manufacturer's instructions. DNA was stored at -20°C, and diluted 10-fold in water before assays. Targeted SNPs in the life history genomic regions were $vgll3_{TOP}$, which has the strongest signal of association with age at maturity in the *vgll3* haploblock on chromosome 25, and $six6_{TOP.LD}$ in the *six6* haploblock on chromosome 9 with complete linkage disequilibrium with the $six6_{TOP}$ SNP reported in (Barson et al. 2015). For *six6*, forward primers were CAGGCTAGGAGCCCAAAAATG[C] and CCAGGCTAGGAGCCCAAAAATG[A] where [A] and [C] represents HEX/FAM labelled allele specific differences in primers corresponding to late and early maturing alleles respectively, and common reverse primer was GTCTGTTGCTTGTGTTTGTGTGTTTACTA. Genotype scores for all markers were called by eye, in which different genotypes formed distinct clusters based on fluorescence intensities of FAM and HEX dyes.

*Food availability treatment*

The treatment started in August 2020, at least three weeks after PIT-tagging for each family and one week after splitting each family to two tanks for low food and high food treatments (Fig. 1). Mean density in the tanks in the beginning of the treatments was 1.29 g $L^{-1}$, N = 145–152 $tank^{-1}$). In the beginning of the experiment, the relative age of the fish was approximately 2050 degree days, or 1313 Tau as calculated with the formula of Gorodilov (1996). Fish in the high food treatment tanks were weighed and measured before the treatment, including 2d fasting and anaesthesia (see above). Fish in the low food treatment were measured once for SMR (as described below, for a separate study) before the treatment, including in total 3d fasting, after which they were weighed to nearest 0.01g, and measured to nearest mm. Densities of fish biomass in the tanks at the end of the treatments were on average 1.8 and 2.8 g L-1 in the low food and high food treatments, respectively. No signs of aggression among the fish were observed during the treatment. Feed rations were adjusted weekly during the treatments.

After two weeks, approximately 25 fish were randomly captured from each tank by netting, sedated with buffered methanesulfonate (100 $mgL^{-1}$) and weighed to recalculate rations based on true biomass (mean mass of fish × nr of fish in the tank). We subsequently balanced the densities of fish in the low-food and high-food tanks for each family by culling fish from the high-food treatment with an overdose of buffered methanesulfonate (approx. 25 fish culled from high food tanks).

Fish of desired genotypes were randomly picked from the rearing tanks for acclimation by netting, apart from the high food treatment from the 4th family where some fish had grown too large for the respirometers. All fish from this tank were anaesthetised, weighed, and measured four days before their respirometry trials, and appropriate size fish (max. length 82mm) were selected for the trials from each genotype (*vgll3* and *six6* genotype frequencies were not different between these two size groups, $Chi^2 = 5.5$, $df = 7$, $p = 0.6$).

*Acclimation conditions before metabolic rate measurements*

Temperatures of the acclimation tanks were regulated to offset the higher temperature of the room (17°C) using a temperature controller (TS1000, H-Tronic Gmbh, Germany) with a temperature probe (PT-1000) located in the acclimation tank, and an Eheim 300 pump (Eheim, Deizisau, Germany) located in a separate cooling tank that was set to 9°C. Water in the cooling tank was kept cool using a Teco 2000 (Teco, Ravenna, Italy) aquarium cooling unit. To refresh water in the system, the cooler tank received a slow (approx. 1L $min^{-1}$) flow of water from the same system as the rearing tanks, which was slow enough for the cooling unit to keep the temperature stable. Constant aeration and water circulation using two Eheim 300 pumps ensured even and well-oxygenated conditions for all fish in the acclimation tanks. Two acclimation tanks were used for two separate batches of fish at once. One 80-cm Juwel Novolux LED-light was placed over each acclimation tank. The photoperiod matched that of the rearing tanks (which mimicked natural day length) and was adjusted weekly.

*Respirometer design*

A 16-channel intermittent-flow respirometer, built in-house, was located next to the acclimation tanks. Tank volume was 164L and temperature was 11°C ± 0.1°C, controlled using an independent cooling system similar to the acclimation tanks, with constant inflow of lake water from the same system as the rearing tanks at approx. 1L $min^{-1}$, which was passed through a 60 µm filter. Chambers for the respirometer were made using borosilicate glass tubing (length 120mm, inner diameter 38mm, wall thickness 3.2mm) (Schott, Finnish Special Glass). The 3D-printed 'HeiBer'-caps were designed by Heidrikur Bergsson, University of Copenhagen (Fig. S2, https://zenodo.org/record/4062429#.YMSW7h1RVTY). The chamber caps were prepared by 3D printing of polyethylene terephthalate glycol (PETG, Devil Design, Mikolow, Poland) using a ZMorph VX 3D printer equipped with a single extruder (ZMorph S.A., Wrocław, Poland). The G-code for the 3D printer was generated with Voxelizer 2 software (ZMorph). The caps were printed using a nozzle size of 0.4 mm, layer height of 0.12 mm, 100% infill, and extruder temperature of 250°C, while the worktable temperature was set to 70°C. For overhanging sections deviating less than 60° from horizontal plane, separate support structures were used. Other parameters were according to the default settings for PETG printing in Voxelizer.

Water inside the chambers was mixed by a plastic disk attached to each cap to distribute the flow. The disks were 3D-printed using the same method as the caps and attached using stainless steel screws. Caps were sealed with nitrile butadiene rubber o-rings and connected to the recirculation and flush pumps with Tygon tubing (Tygon S3 E-3603, Saint-Gobain, Paris, France). The recirculation system was verified as waterproof by filling it with water after plugging the flush pump and probe connections. For flushing the chambers, one end of tubing from each chamber was placed above the water surface. Water was recirculated in chambers using 16 submersible 3-6V DC pumps (amphibious type, unbranded/generic, China). Chambers were flushed using four magnetic 5-12V DC (DC30C, ANSELF, China), from each of which the flow was directed to four chambers at approximately equal rates. Flush- and recirculation pumps were controlled using PumpResp controllers (4-channel model, FishResp, Finland, https://fishresp.org/pumpresp). The information about time and corresponding phase (i.e., flush or measurement) was recorded by the PumpResp controllers to a PC. Mean flow rate within the chamber was calculated based on pump outflow rate and chamber diameter from 5 pumps (mean ± SD: closed phase 0.3 ± 0.1 cm $s^{-1}$ and flush phase 0.9 ± 0.1 cm $s^{-1}$ during SMR; and closed phase 1.5 ± 0.2 cm $s^{-1}$ during MMR). Flow had no apparent effect on fish movements in the chambers in neither SMR nor MMR measurements. Flow of all recirculation pumps was calibrated daily, separately for SMR and MMR measurements. The ratio of water volume in the chamber and tubing to fish body mass was on average 37 (range 18–82, fish body mass was 1.49–6.89 g).

Oxygen saturation (mg $O_2$ $L^{-1}$) was measured at 0.5Hz in the recirculation loops using optical Robust Oxygen sensors connected to Firesting Oxygen meters (Pyroscience, Gmbh, Aachen, Germany). The sensors were connected to tubing between the recirculation pump and the chamber using a plastic T-piece and a small piece of silicon tubing. All 16 sensors were calibrated simultaneously with $O_2$-free water, made using sodium sulphate, and with air-saturated water once before the first measurements. Temperature-compensation for $O_2$ saturation was based on TSUB21-CL5 (Pt100) temperature sensors (Pyroscience, Gmbh, Aachen, Germany), which were placed in the middle of the respirometer tank and connected to each of the Firesting meters.

*Standard metabolic rate measurements*

Before the SMR measurements, background bacterial respiration was measured in empty chambers for one measurement cycle (5 min flush, 2 min wait period, 13 min measurement). Thereafter fish were caught into plastic cups with water (to limit exposure to air) and transferred to the measurement chambers. Measurements were started immediately using the same cycle length as the background respiration. After a few minutes the measuring tank was covered with a light-blocking canvas, and the rest of the measurement was completed in the dark. Background oxygen consumption was measured for one cycle again after the fish were removed from the chambers to account for potential increase in bacterial respiration during the trial. After each family was measured (5d of measurements), the respirometer was cleaned with 10% bleach and all chamber parts scrubbed to limit bacterial growth, followed by thorough rinsing with water.

*Maximum metabolic rate measurements*

The fish were removed from the chambers using plastic cups, their PIT-tags recorded, and they were kept in 10-L buckets at 11°C with aeration (8 fish in each bucket) until the chase tests. During the chase, individual fish were encouraged to swim by hand in a rectangular container (length 48 x width 27 cm) with 8L water (at a similar temperature as acclimation tanks, range 10.6–11.3°C). Chasing was continued for 3 min, at which point all fish had completely fatigued, as they were easy to handle and swam very slowly if at all. The fish were then rapidly (within 20s) transferred to the respirometry chambers using cups (the chambers were flushed, and the measurements started before the fish entered the chambers). No air exposure was used, avoiding the risk of gill lamellae collapse (Cook *et al.* 2015), hence making the MMR measurement more ecologically relevant. Two people chased one fish each at once, and the 16 individuals from each batch were processed within 2h. Water in the chase container was replaced after each individual. After at least one MMR measurement cycle (minimum cycle length included in analysis 5.5min), fish were removed from the chambers, their PIT-tags read, and they were placed back into 10-L buckets with aeration at 11°C before they were euthanised the same day. After all fish in one batch were measured for MMR, background respiration was immediately measured for one cycle in the empty chambers. This background respiration measurement also served as the pre-measurement background measurement for the next batch of SMR that started within approx. 1h.

*Standard metabolic rate data analysis*

The final minute of each measurement phase was trimmed prior to analysis to compensate for minor variation in timing in the start of the flush phase, therefore the duration of each measurement phase was 12 min. Background respiration was accounted for using the R package *FishResp* (Morozov *et al.* 2019), assuming linear growth of microorganisms inside a respirometry chamber over time (only weak background respiration was observed: mean ± SD of background respiration compared to total respiration was 2.4 ± 1.7 %). Water in the respirometer was hypoxic only very briefly during the closed phases of SMR measurements ($O_2$ saturation was >8.2 mg $L^{-1}$ 99% of the time and <6.5 mg $L^{-1}$ < 0.009% of the time). As a result of frequent measurement of oxygen during the measurement phase (every 2s) and the large number of measurements (an average of 60 measurement phases for each individual), identifying non-linear slopes based on $R^2$ values and visual assessment alone was ineffective (see also: Chabot et al. 2021). We therefore applied three complementary approaches: 1) we excluded measurements where $R^2$ of the slope $O_2$ level vs. time < 0.95, which removed slopes with clear effects of fish activity or pump malfunction. 2) We excluded 'outlier' datapoints that showed a large deviation from the fitted regression line of $O_2$ mg $L^{-1}$ vs. time. Outliers were defined as absolute residual values larger than 0.5 (mg O2 $L^{-1}$), and the entire measurement phase was excluded from the analysis if >10% of the datapoints in a slope were flagged as outliers. 3) We evaluated the curvature of the slopes and excluded the measurement if the derivative of the quadratic fit (collected every 2min during the measurement) had a coefficient of variation > 15%. This approach very closely matched a manually conducted evaluation of linearity (Fig. S4). Finally, all accepted slopes were checked visually to verify linearity. Individuals were retained in the analysis if they had more than 20 slopes remaining after the filtering. The filtering excluded five fish from the low food treatment (with 4–19 slopes) and one fish from the high food treatment (with 14 slopes). Next, based on visually identifying abnormal data across measurements collected from each fish (caused, e.g., by a too slow recirculation) and on observing large (>6mm) air bubbles in the chambers after the measurement, we further excluded five individuals from the low food group and two from the high food group from analysis. Small air bubbles (<6mm) in six instances were corrected for by reducing the total volume of the chamber by the estimated volume of the bubble (which had a negligible effect on the results). Mass-adjusted MLND-SMR values were highly correlated with mass-adjusted SMR calculated using quantile methods (Pearson-r 0.94 and 0.95 for q=0.1 and q=0.2, respectively, both $p < 0.001$).

*Maximum metabolic rate data analysis*

We excluded the first 20s after each fish entered the chamber from the analysis, as these data were noisy from water mixing and, in some cases, from releasing air bubbles from the chamber. We excluded 2 individuals from MMR measurements and a further 12 individuals from both SMR and MMR analysis due to human error in the measurements. The moment of fish entering the chamber was determined from the oxygen traces using a custom-made function that identified the change in slope as the chamber was closed (see data availability). The metabolic rate of fish decreased with time after the chase, i.e., the absolute slope of $O_2$ level vs. time decreased with time since the end of the chase. First, we used package *respR* (Harianto, Carey & Byrne 2019) with the function *auto_rate*, fitting one and two-minute windows as suggested by (Little *et al.* 2020) (example slope in Fig. S5A). For each window width, MMR was selected as the maximum slope with $R^2$ >0.95, which omitted slopes with low accuracy from the analysis (e.g., when the slope included outlier points or pump malfunction). The 1-min window size was strongly correlated with, but gave higher MMR values than, those obtained using 2-min windows for the analysis (means: 1-min = 682 mg $O_2$ $L^{-1}$ $h^{-1}$ and 2-min = 630 mg $O_2$ $L^{-1}$ $h^{-1}$). In the spline method, the slope for MMR was obtained from the beginning of the accepted measurement period for each fish.

### 2. Figures

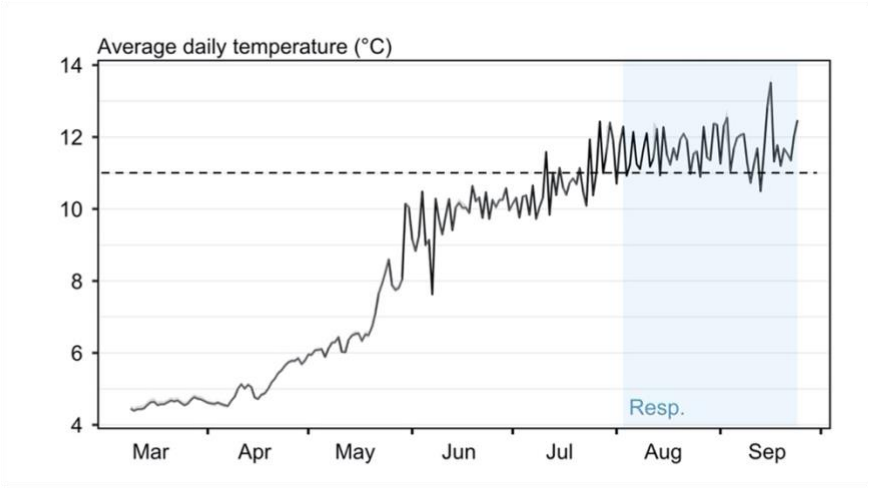

Fig S1. Daily temperature during fish rearing and the experimental period. Minimum and maximum temperature across tanks shown in the (very narrow) grey area. The timing of the respirometry experiment is shaded in blue – note that fish were acclimated to a stable temperature for two days before respirometry, which was performed at the same temperature (11°C, dashed line).

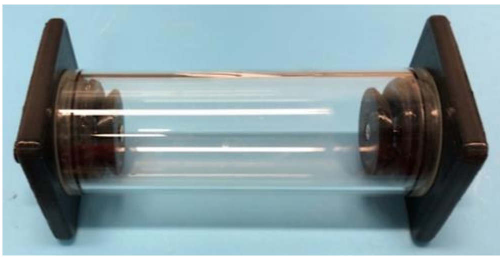

Fig S2. Photo of the respirometry chamber made of 3D-printed caps and a glass tube.

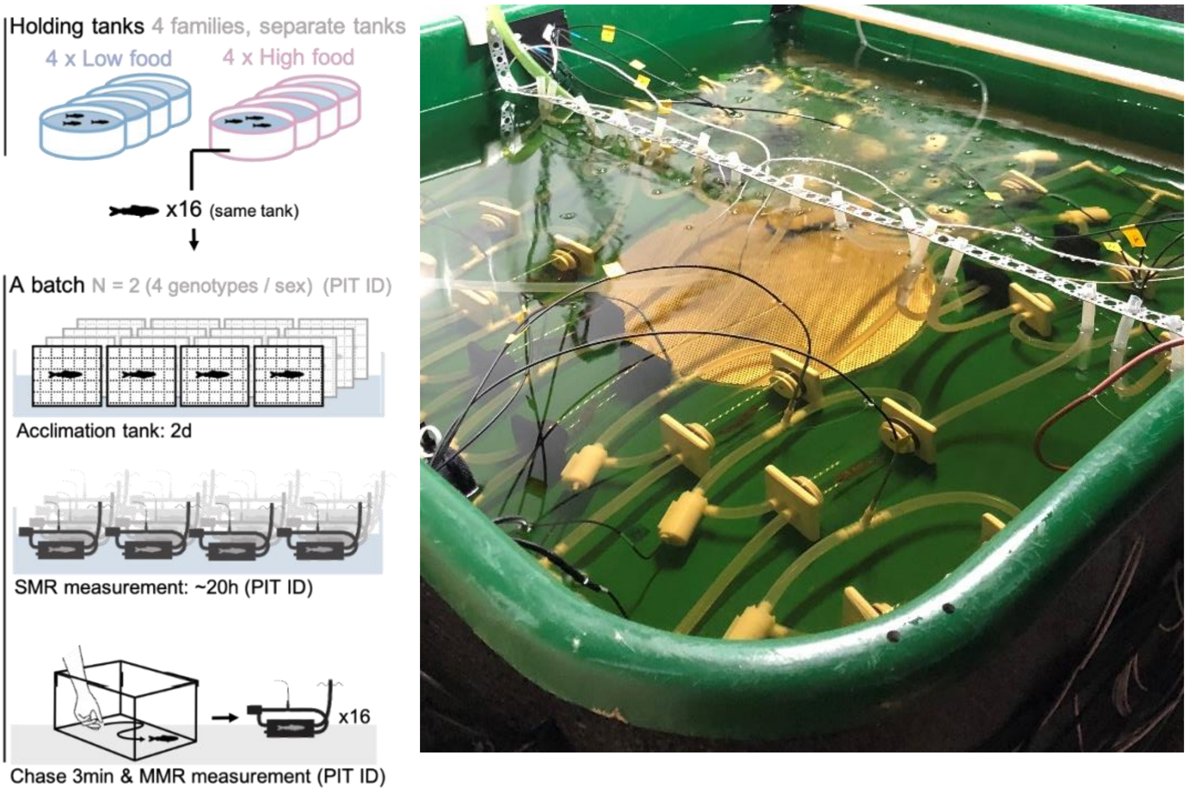


Fig S3. Diagram showing the procedure of acclimation, followed by SMR and MMR measurements (left). Fish were identified using PIT IDs at different steps of the study. Photo of fish in the respirometer (right).

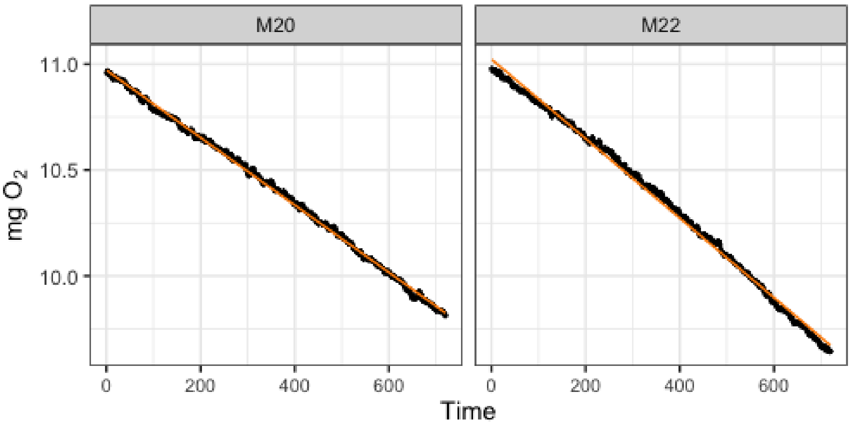


Fig. S4. Examples of linear (left, coefficient of variation 6.9%) and curved (right, coefficient of variation 15.4%) slopes. Linear slope shown with orange line. Both slopes had $R^2$ >0.95.

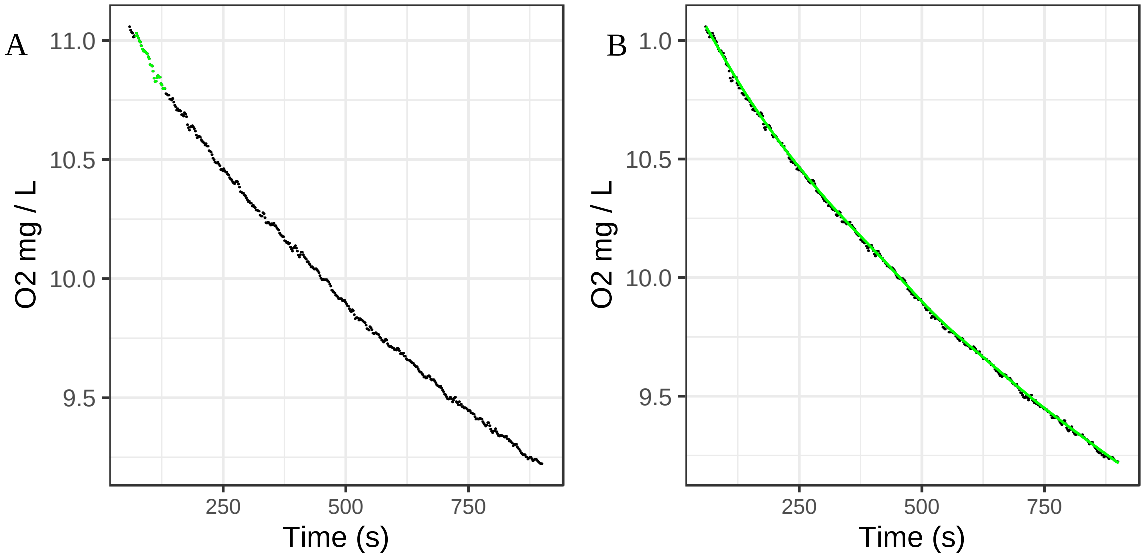


Fig S5. A) An example slope from an MMR measurement with points for the steepest slope identified using a 1-min sliding window using package *respR* (*b* = -0.00414). B) The same slope fitted with a polynomial line in green for the 'spline-MMR' method. Slope was taken from the tangent from the beginning of the fitted line (Time = 1s, *b* = -0.00359). See data availability for raw data and R scripts.

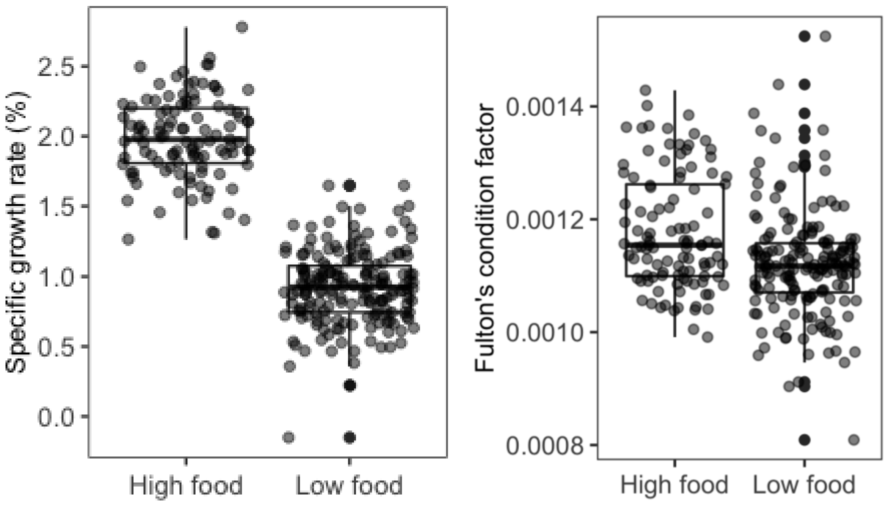


Fig. S6. a) Specific growth rate during and b) Fulton's condition factor in juvenile salmon after food treatments.  N = 107 (high food) and 180 (low food). Points represent individuals. Growth rate was calculated as specific (a.k.a. instantaneous) growth rate as follows: [(ln mass at $t_2$ - ln body mass at $t_1$) / days between $t_1$ and $t_2$)] × 100, where $t$ is date. Fulton's condition factor was calculated as (body mass / length$^3$) × 100.

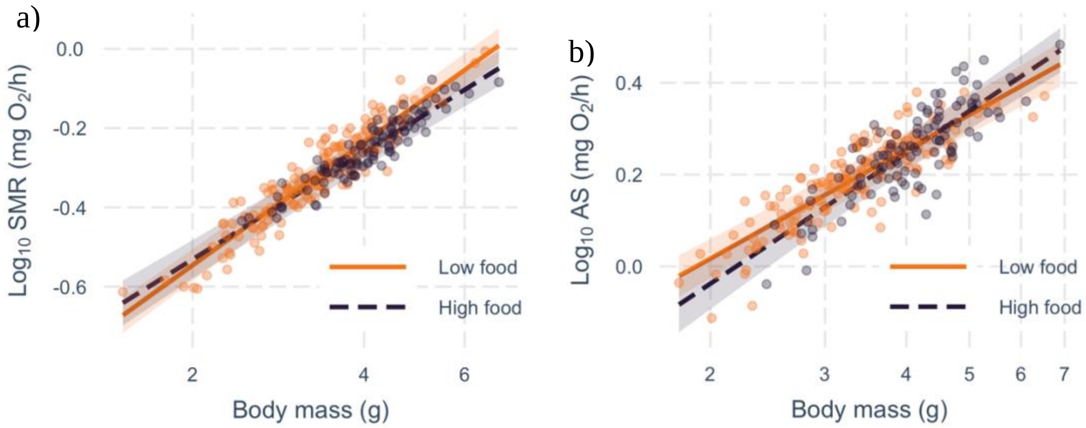


Fig. S7. Partial residuals of a) SMR and b) aerobic scope (AS) with body mass ($\log_{10}$-scale) in high and low food treatments. Shaded areas show 95% prediction intervals for each regression line. N = 104/100 (SMR/AS, high food) and 160/153 (low food). Points represent individuals.

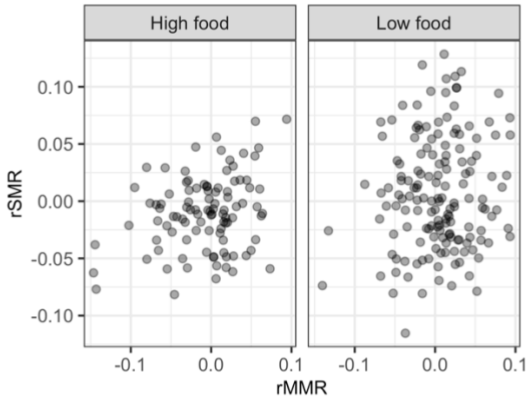


Fig S8. Scatter plots of family- and mass-corrected SMR and MMR in high and low food treatments. Points represent individuals. Correlation coefficients are shown in Table 2. N = 100 (high food) and 153 (low food).

### 3. Tables

Table S1. Information on parental fish and eggs, when available (weights not collected for males). All parental fish were from the same cohort and reared in captivity (crossed in 2011 using fish caught from the wild). Egg weight was calculated as a mean from 20 eggs.

| Family nr | Family ID | Female length (cm) | Female weight (g) | Male length (cm) | Mean egg weight (g) |
|---|---|---|---|---|---|
| 1 | F1 | 79.3 | 8355 | 64.7 | 0.12 |
| 2 | F3 | 74.5 | 5700 | 69.6 | 0.15 |
| 3 | F4 | n.a. | n.a. | 69.8 | 0.12 |
| 4 | F8 | 74.7 | 5600 | 72 | 0.14 |

Table S2. The total number of homozygous individuals in each genotype-sex combination across families and treatments. Genotypes for *vgll3* and *six6*: EE (homozygous for early maturation allele), LL (homozygous for late maturation allele). Total N = 107 and 183 for high food and low food, respectively.

| | | vgll3*EE/ six6*EE | | vgll3*EE/ six6*LL | | vgll3*LL/ six6*EE | | vgll3*LL/ six6*LL | |
|---|---|---|---|---|---|---|---|---|---|
| | | F | M | F | M | F | M | F | M |
| High food | Family 1 | 0 | 0 | 0 | 0 | 3 | 0 | 4 | 4 |
| | Family 2 | 4 | 5 | 4 | 4 | 4 | 3 | 5 | 3 |
| | Family 3 | 2 | 6 | 4 | 4 | 4 | 4 | 4 | 4 |
| | Family 4 | 5 | 3 | 4 | 5 | 6 | 2 | 3 | 4 |
| | Total | 11 | 14 | 12 | 13 | 17 | 9 | 16 | 15 |
| Low food | Family 1 | 4 | 6 | 4 | 7 | 6 | 2 | 4 | 4 |
| | Family 2 | 7 | 6 | 5 | 6 | 6 | 6 | 6 | 6 |
| | Family 3 | 5 | 7 | 6 | 7 | 6 | 5 | 6 | 6 |
| | Family 4 | 6 | 6 | 6 | 6 | 6 | 6 | 6 | 6 |
| | Total | 22 | 25 | 21 | 26 | 24 | 19 | 22 | 22 |

Table S3. Definitions of model parameters. All fixed effects were centred for the analysis (mean of 0).

| **Fixed effects** | **Explanation** |
|---|---|
| Vgll3 genotype | Homozygous EE / -0.5 or LL/0.5 |
| Six6 genotype | Homozygous EE / -0.5 or LL/0.5 |
| Sex | M / 0.5 = male, F /-0.5 = female |
| Treatment | High food (fed to satiation daily) (-0.5) /<br>Low food (fed to satiation 2x $wk^{-1}$) (0.5) |
| Log10 body mass | Log10-transformed body mass (g) measured after SMR and MMR trials. Scaled and centred for analysis. |
| Test order | Covariate, 1–8, indicating the order of pairs of fish tested each day for MMR. |
| **Random effects** | |
| Family | Family + Tank effect (each family in separate tank for each treatment) |
| Batch | The 16 individuals used in respirometry each day |
| Chamber.No | The chamber in which MMR was measured after chase (CH1–CH16) |
| Initial | Initial of the person performing chase test |

Table S4. Top-ranked models used in model averaging. Random effects were as in the full model in all models.

| | | Terms in the model | AICc | delta-AICc |
|---|---|---|---|---|
| SMR | Mod1 | Six6 + Vgll3 + log10(Mass)) + Treatment + Six6:Vgll3 + Six6:Treatment + log10(Mass)):Treatment | -916.63 | 0.00 |
| | Mod2 | Six6 + log10(Mass)) + Treatment + Six6:Treatment + log10(Mass)):Treatment | -915.96 | 0.67 |
| | Mod3 | Six6 + Vgll3 + log10(Mass)) + Treatment + Six6:Treatment + log10(Mass)):Treatment | -915.94 | 0.70 |
| | Mod4 | log10(Mass)) + Treatment + log10(Mass)):Treatment | -915.87 | 0.76 |
| | Mod5 | Vgll3 + log10(Mass)) + Treatment + log10(Mass)):Treatment | -915.57 | 1.06 |
| | Mod6 | Six6 + Vgll3 + log10(Mass)) + Treatment + Six6:Vgll3 + Six6:Treatment + Vgll3:Treatment + log10(Mass)):Treatment | -914.87 | 1.77 |
| | Mod7 | Six6 + Vgll3 + log10(Mass)) + Treatment + Six6:Vgll3 + Six6:Treatment | -914.83 | 1.81 |
| | Mod8 | Six6 + Vgll3 + log10(Mass)) + Treatment + Six6:Vgll3 + log10(Mass)):Treatment | -914.77 | 1.87 |
| | Mod9 | Six6 + log10(Mass)) + Treatment + log10(Mass)):Treatment | -914.65 | 1.99 |
| MMR | Mod1 | Six6 + Vgll3 + log10(Mass)) + Six6:Vgll3 | -883.45 | 0.00 |
| | Mod2 | Six6 + Vgll3 + log10(Mass)) + Treatment + Six6:Vgll3 | -882.53 | 0.92 |
| | Mod3 | Six6 + Vgll3 + log10(Mass)) + Order) + Six6:Vgll3 | -882.18 | 1.27 |
| | Mod4 | Six6 + Vgll3 + log10(Mass)) + Treatment + Six6:Vgll3 + log10(Mass)):Treatment | -881.88 | 1.58 |
| | Mod5 | Six6 + Vgll3 + log10(Mass)) + Sex + Six6:Vgll3 + Vgll3:Sex | -881.81 | 1.64 |
| | Mod6 | Six6 + Vgll3 + log10(Mass)) + Sex + Six6:Vgll3 | -881.60 | 1.85 |
| AS | Mod1 | Vgll3 + log10(Mass)) + Treatment + log10(Mass)):Treatment | -721.95 | 0.00 |
| | Mod2 | Six6 + Vgll3 + log10(Mass)) + Treatment + log10(Mass)):Treatment | -720.89 | 1.06 |
| | Mod3 | Vgll3 + log10(Mass)) + (1 \| initial) | -720.61 | 1.34 |
| | Mod4 | Vgll3 + log10(Mass)) + Treatment + Vgll3:Treatment + log10(Mass)):Treatment | -720.61 | 1.34 |
| | Mod5 | Six6 + Vgll3 + log10(Mass)) + Treatment + Six6:Vgll3 + log10(Mass)):Treatment | -720.59 | 1.36 |
| | Mod6 | Vgll3 + log10(Mass)) + Sex + Treatment + Vgll3:Sex + log10(Mass)):Treatment | -720.49 | 1.46 |
| | Mod7 | Vgll3 + log10(Mass)) + Sex + Treatment + log10(Mass)):Treatment | -720.43 | 1.52 |
| | Mod8 | Vgll3 + log10(Mass)) + Treatment + (1 \| initial) | -720.34 | 1.61 |
| | Mod9 | Vgll3 + log10(Mass)) + Order) + Treatment + log10(Mass)):Treatment | -720.06 | 1.89 |

Table S5. Averaged parsimonious models for log10 metabolic phenotypes. All variables were centred to a mean of 0 (the category with a positive value is shown in parentheses), log10 body mass was scaled and centred. Significant effects (p < 0.05) shown in bold. Importance indicates the Sum of Akaike weights over all models including the explanatory variable.

| | Coefficient | Estimate | SE | z | Importance | P |
|---|---|---|---|---|---|---|
| SMR | **Log10 BM** | **0.1025** | **0.003** | **31.846** | **1.00** | **<0.001** |
| | Treatment (LF) | 0.0202 | 0.012 | 1.725 | 1.00 | 0.09 |
| | Log10 BM:Treatment (LF) | 0.0125 | 0.007 | 1.753 | 0.92 | 0.08 |
| | *Six6* (LL) | -0.0021 | 0.004 | 0.464 | 0.76 | 0.64 |
| | *Vgll3* (LL) | -0.0047 | 0.005 | 0.907 | 0.67 | 0.37 |
| | *Six6* (LL):Treatment (LF) | -0.012 | 0.012 | 0.978 | 0.61 | 0.33 |
| | *Six6* (LL):*Vgll3* (LL) | -0.0067 | 0.01 | 0.668 | 0.42 | 0.50 |
| | *Vgll3* (LL):Treatment (LF) | 0.0005 | 0.003 | 0.155 | 0.08 | 0.88 |
| MMR | *Six6* (LL) | -0.0073 | 0.005 | 1.38 | 1.00 | 0.168 |
| | ***Vgll3* (LL)** | **-0.0179** | **0.005** | **3.412** | **1.00** | **0.001** |
| | **Log10 BM** | **0.087** | **0.003** | **28.282** | **1.00** | **<0.001** |
| | ***Six6* (LL):*Vgll3* (LL)** | **-0.0228** | **0.01** | **2.17** | **1.00** | **0.03** |
| | Treatment (LF) | 0.0023 | 0.005 | 0.481 | 0.32 | 0.63 |
| | Test order | -0.0006 | 0.002 | 0.279 | 0.15 | 0.78 |
| | Log10 BM:Treatment (LF) | -0.0011 | 0.004 | 0.294 | 0.13 | 0.77 |
| | Sex (male) | -0.0007 | 0.003 | 0.238 | 0.24 | 0.81 |
| | *Vgll3* (LL):Sex (male) | 0.002 | 0.007 | 0.314 | 0.13 | 0.75 |
| AS | ***Vgll3* (LL)** | **-0.0176** | **0.007** | **2.56** | **1.00** | **0.01** |
| | **Log10 BM** | **0.0899** | **0.004** | **20.331** | **1.00** | **<0.001** |
| | Treatment (LF) | 0.0123 | 0.009 | 1.419 | 0.90 | 0.16 |
| | Log10 BM:Treatment (LF) | -0.0136 | 0.01 | 1.328 | 0.80 | 0.18 |
| | *Six6* (LL) | -0.0016 | 0.004 | 0.365 | 0.22 | 0.72 |
| | *Vgll3* (LL):Treatment (LF) | -0.0013 | 0.006 | 0.222 | 0.10 | 0.82 |
| | *Six6* (LL):*Vgll3* (LL) | -0.0019 | 0.007 | 0.269 | 0.10 | 0.79 |
| | Sex (male) | -0.001 | 0.004 | 0.283 | 0.19 | 0.78 |
| | *Vgll3* (LL):Sex (male) | 0.002 | 0.007 | 0.27 | 0.10 | 0.79 |
| | Test order | -0.0002 | 0.002 | 0.132 | 0.08 | 0.90 |

BM = Body mass, LF = Low food

Table S6. The poportions of variance of mass-adjusted MMR and AS explained by genotype effects that were statistically significant (Table 1).

| | Term | Estimate | 2.5% C.I. | 97.5% C.I. | Df |
|---|---|---|---|---|---|
| MMR | All genotype effects | 0.0501 | 0.0209 | 0.1188 | 4 |
| | Vgll3 | 0.0323 | 0.0006 | 0.0984 | 3 |
| | Vgll3 x Six6 | 0.0130 | 0.0000 | 0.0770 | 3 |
| | Vgll3 + Vgll3 x Six6 | 0.0463 | 0.0166 | 0.1143 | 2 |
| AS | Vgll3 | 0.018 | 0.001 | 0.064 | 1 |

Table S7. Linear mixed model for log10 absolute aerobic scope including mass-adjusted standard metabolic rate (rSMR) as covariate. Significant effects shown in bold.

| Coefficient | Estimate | SE | SSq | Den DF | F | p |
|---|---|---|---|---|---|---|
| Intercept | 0.211 | 0.017 | | | | |
| Treatment (LF) | 0.013 | 0.008 | 0.008 | 239.211 | 2.837 | 0.093 |
| Sex (male) | -0.004 | 0.007 | 0.001 | 242.967 | 0.442 | 0.507 |
| ***Vgll3* (LL)** | **-0.018** | **0.007** | **0.017** | **241.785** | **6.428** | **0.012** |
| *Six6* (LL) | -0.009 | 0.007 | 0.004 | 242.492 | 1.593 | 0.208 |
| **Log10 BM** | **0.090** | **0.004** | **1.178** | **247.148** | **437.872** | **<0.0001** |
| Test order | -0.003 | 0.005 | 0.001 | 14.473 | 0.344 | 0.567 |
| rSMR | -0.007 | 0.004 | 0.006 | 236.883 | 2.275 | 0.133 |
| Treatment (LF):log10 BM | -0.017 | 0.009 | 0.010 | 245.213 | 3.733 | 0.054 |
| Treatment (LF):*Vgll3* (LL) | -0.015 | 0.014 | 0.003 | 242.507 | 1.068 | 0.302 |
| Treatment (LF):Six6 (LL) | -0.001 | 0.014 | 0.000 | 243.856 | 0.005 | 0.945 |
| Sex (male):*Vgll3* (LL) | 0.024 | 0.014 | 0.008 | 239.953 | 3.052 | 0.082 |
| Sex (male):*Six6* (LL) | 0.010 | 0.014 | 0.002 | 239.993 | 0.603 | 0.438 |
| *Vgll3* (LL):*Six6* (LL) | -0.022 | 0.014 | 0.007 | 244.057 | 2.722 | 0.100 |

BM = Body mass, LF = Low food
